## Supplementary figures and images for "The spindle assembly checkpoint and the spatial activation of Polo kinase determine the duration of cell division and prevent neural stem cells tumor formation"

### Figure S1

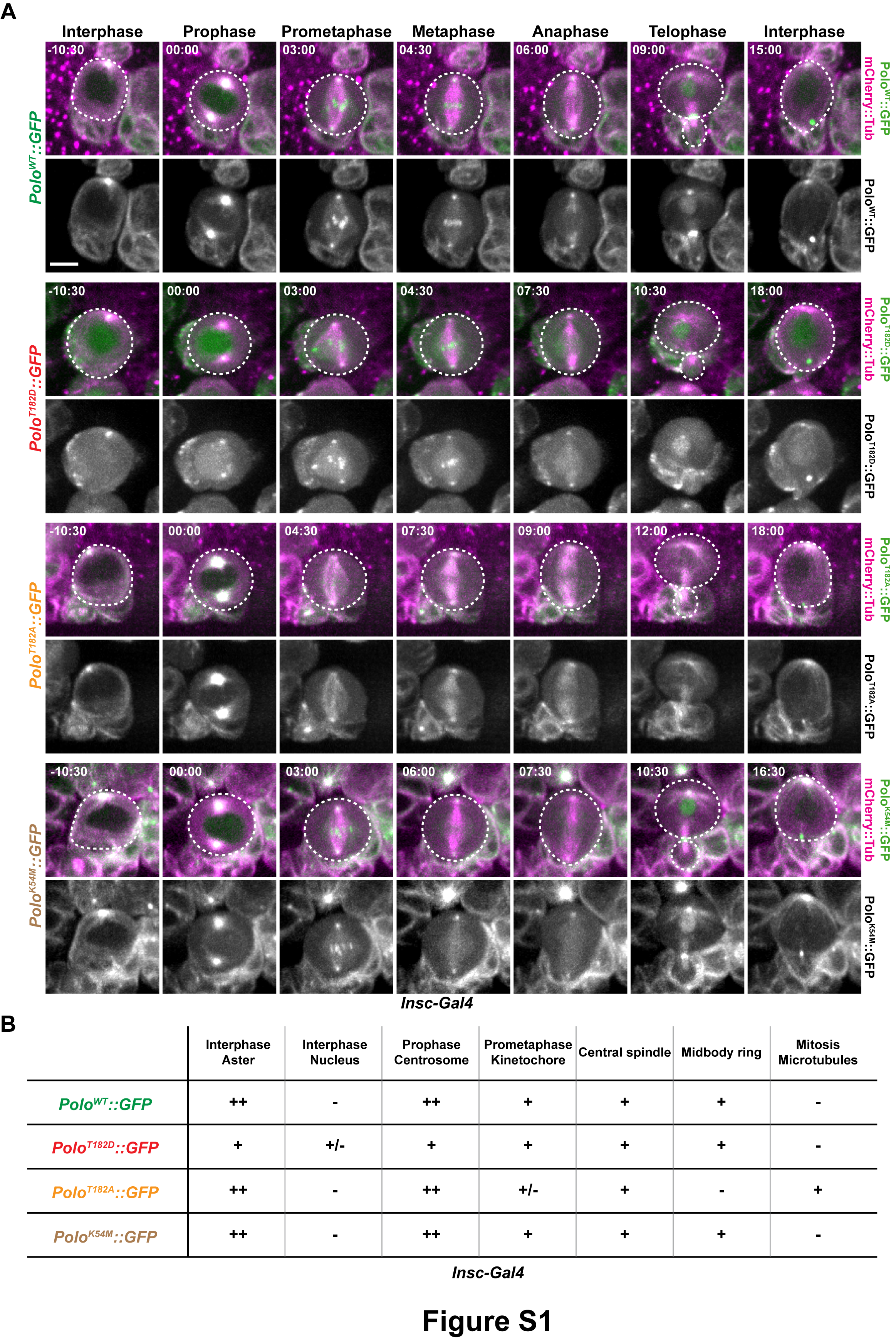

### Figure S2

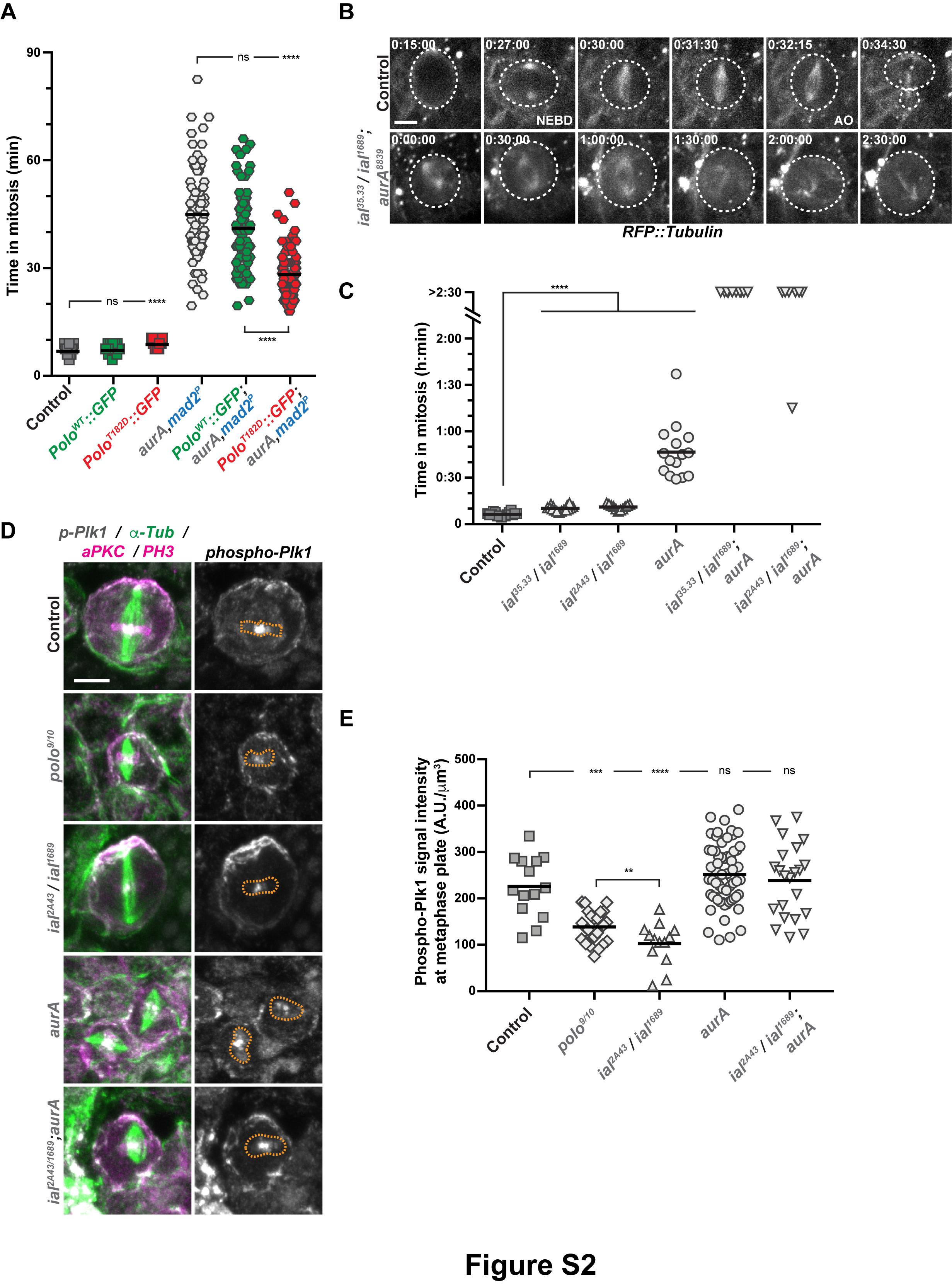

### Figure S3

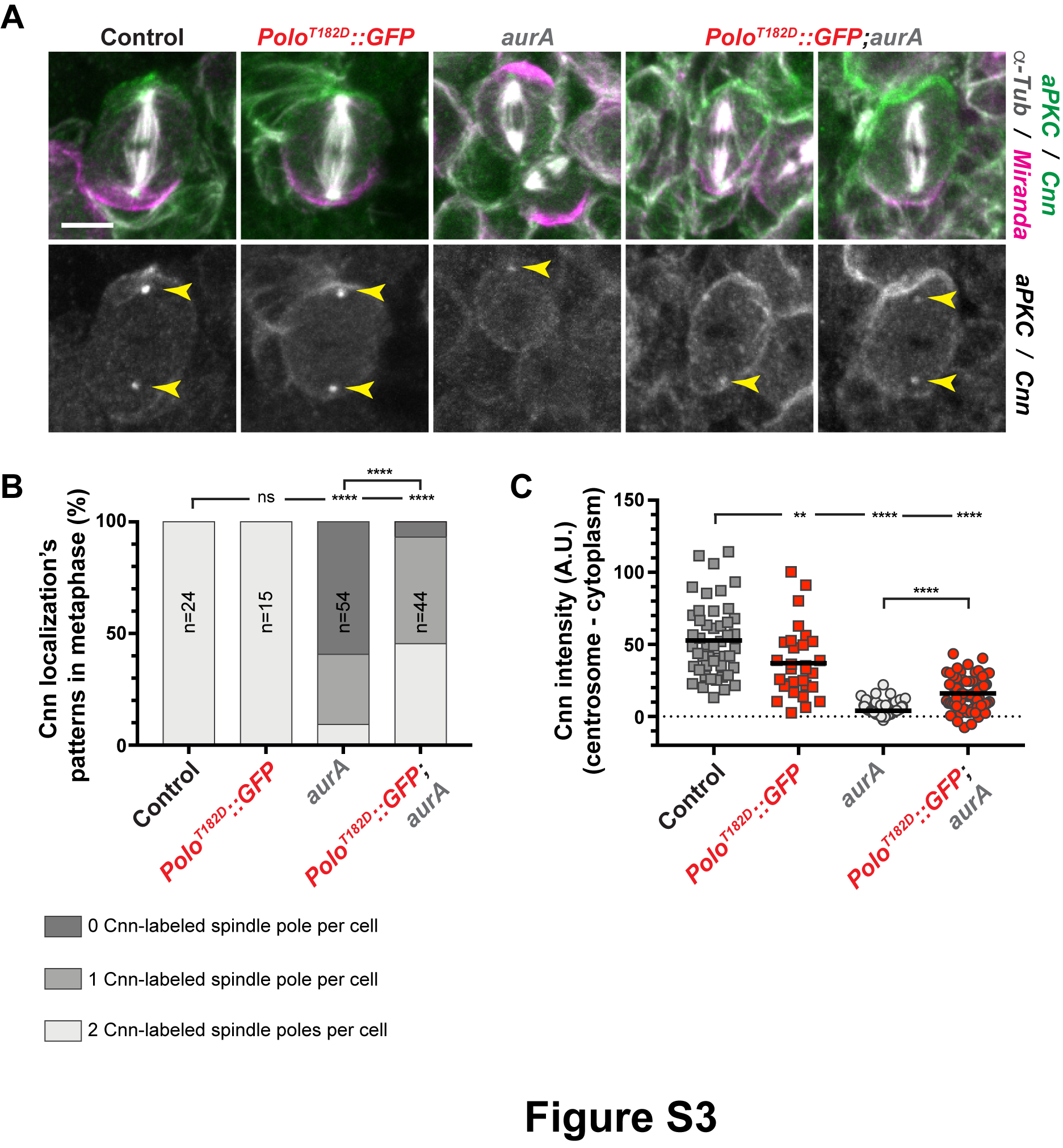
